# Sphingosine-1-phosphate signaling inhibition suppresses Th1-like Treg generation by reversing mitochondrial uncoupling

**DOI:** 10.1101/2024.05.20.594966

**Authors:** Rachel Coulombeau, Claudia Selck, Nicolas Giang, Abdulrahman Al-Mohammad, Natalie Ng, Allison K Maher, Rafael Argüello, Antonio Scalfari, James Varley, Richard Nicholas, Margarita Dominguez-Villar

## Abstract

Inflammatory environments induce the generation of dysfunctional IFN_γ_^+^T-bet^+^FOXP3^+^ Th1-like Tregs, which show defective function and are found in autoimmune conditions including multiple sclerosis (MS). The pathways that control the generation of Th1-like Tregs are not well understood. Sphingosine-1-phosphate (S1P) signaling molecules are upregulated in Th1-like Tregs, and *in vivo* S1P inhibition with Fingolimod (FTY720) inhibits the expression of genes responsible for Treg plasticity in MS patients. However, the underlying mechanisms are unknown. Here we show that S1P signaling inhibition by FTY720 inhibits the generation of Th1-like Tregs and rescues their suppressive function. These effects are mediated by a decrease in mTORC1 signaling and reversal of the mitochondrial uncoupling that Tregs undergo during their reprogramming into Th1-like Tregs *in vitro*. Finally, these results are validated in *in vivo* generated Th1-like Tregs, as the Tregs from MS patients treated with FTY720 display decreased Th1-like Treg frequency, increased suppressive function, and mitochondrial metabolism rebalance. These results highlight the involvement of mitochondrial uncoupling in Treg reprogramming and identify S1P signaling inhibition as a target to suppress the generation of dysfunctional Th1-like Tregs.

---

Regulatory T cell (Treg) plasticity occurs as a physiological adaptation to the changing environment and the sensing of danger signals, which allows Tregs to control specific helper T cell responses^1^. However, aberrant Treg acquisition of an effector phenotype with secretion of pro-inflammatory cytokines is characteristic of several autoimmune conditions including multiple sclerosis (MS)^2, 3^, type 1 diabetes^4^, rheumatoid arthritis^5^, systemic sclerosis^6^ and autoimmune hepatitis^7^. MS is an autoimmune disease of the central nervous system (CNS) characterized by infiltration of activated inflammatory cells into the CNS that damage myelin and axons^8^. In MS, Tregs display a Th1-like phenotype characterized by the increased secretion of the pro-inflammatory cytokine IFN_γ_, upregulation of T-bet and decreased suppressive capacity^3^. The main known pathway involved in the acquisition of a Th1-like phenotype by Tregs is the PI3K/AKT/FOXO signaling^2, 9, 10^, but multiple soluble factors, including IL-12, NaCl and oleic acid^3, 11, 12^ and cell surface receptors such as TIGIT and neuropilin-1^13, 14^ have been shown to either induce or inhibit Treg plasticity towards a Th1-like phenotype, many of them by feeding into the PI3K/AKT pathway.

Tregs exhibit a unique metabolic profile, relying heavily on lipid metabolism and oxidative phosphorylation for their function^15, 16^ and less on glycolysis^17^ compared to effector T cells. In support of this, FOXP3 expression has been shown to promote mitochondrial respiration^18^, suggesting that proper Treg function requires mitochondrial integrity^19, 20^. For example, mice with a Treg-specific deletion of complex III of the mitochondrial respiratory chain develop fatal autoimmunity early in life, without affectation of Treg numbers or FOXP3 expression^21^.

Sphingosine-1-phosphate (S1P) is a bioactive lipid mediator secreted to the extracellular space mostly by vascular endothelial cells, red blood cells and platelets, that is involved in many physiological processes including cell migration, differentiation and survival^22^. S1P signals via five G protein-coupled receptors, i.e. S1PR1-S1PR5, which are expressed on the surface of lymphocytes among other cell types^23^. In T cells, S1P signaling mediates the egress of lymphocytes from secondary lymphoid organs to the target tissue via an S1P gradient, where S1P concentrations are low in lymphoid organs but high in blood and lymph^24^. Thus, inhibition of inflammatory T cell migration from the periphery to the CNS by blocking S1P signaling is a successful therapeutic approach in MS^25, 26^. FTY720 (Fingolimod) is a S1P receptor modulator that is phosphorylated in target cells by sphingosine kinases to become biologically active. Once phosphorylated, FTY720 becomes a high affinity ligand for four S1P receptors (all except S1PR2), acting as a functional antagonist and thereby inhibiting lymphocyte egress from secondary lymph nodes^27^. However, increasing number of works suggest that S1P signaling participates in many additional processes besides cell migration^23^ including dendritic cell and macrophage function, promotion of T cell differentiation to Th1 and Th17 cells and germinal center B cell survival^28, 29, 30, 31^. In this respect, we have previously demonstrated that Treg cells isolated from FTY720-treated relapsing-remitting (RR) MS patients show a decrease in the expression of genes associated with the Th1-like phenotype *ex vivo* compared to baseline, suggesting that FTY720 affects Treg stability independently of cell migration^32^. However, the underlying mechanisms are unknown.

Here we investigated the role of S1P signaling on the regulation of dysfunctional Th1-like Tregs by utilizing FTY720 as a pathway inhibitor. We found that Th1-like Treg generation is suppressed by S1P signaling inhibition by targeting mTORC1 activation. Moreover, Th1-like Tregs undergo mitochondrial uncoupling, with a decrease in mitochondrial dependence and oxygen consumption rate but maintained mitochondrial membrane potential as compared to control Tregs, which is reversed by FTY720 treatment. Finally, by using MS Tregs as a model of *in vivo*-generated Th1-like Tregs, we demonstrate that *in vivo* FTY720 treatment reverses the mitochondrial dysfunction observed in Tregs from untreated RRMS patients. These results underscore the involvement of mitochondrial uncoupling in Treg plasticity and the role of S1P signaling inhibition in restoring mitochondrial function, suggesting that targeting mitochondrial uncoupling is a potential strategy to control Treg stability.

## Results

### S1P signaling inhibition suppresses Th1-like Treg phenotype and function

S1P signaling inhibition by *in vivo* FTY720 (Fingolimod) treatment in RRMS patients leads to a decrease in *IFNG* and *TBX21* gene expression^32^ and increased expression of TIM-3, a marker of highly suppressive Tregs^33^, by Tregs. As Tregs express S1P receptors^29, 34^, we decided to examine whether inhibition of the Th1-like Treg phenotype observed *in vivo* was due to a direct S1P signaling inhibition on Treg cells. For this, we stimulated Tregs isolated from healthy individuals in the presence or absence of IL-12, used as a model to induce Th1-like Tregs *in vitro*^3, 35^, and we tested various concentrations of FTY720 for their effect in regulating Th1-like Treg markers (Fig. 1). FTY720 did not cause any significant increase in cell death as compared to vehicle-treated Tregs at any of the concentrations used (Suppl. Fig. 1). While no effects of FTY720 were observed in the expression of *IFNG, IL10, TBX21* or *FOXP3* in vehicle-treated Tregs, FTY720 significantly decreased the expression of *IFNG, TBX21* and *IL10* in Th1-like Tregs. No changes were observed in the expression of *FOXP3* (Fig. 1A). These results were confirmed at protein level, with increasing doses of FTY720 significantly inhibiting the production of IFN_γ_ (Fig. 1B, 1C) while not affecting the expression of FOXP3 (Fig. 1B and Suppl Fig. 2). Moreover, T-bet expression was significantly reduced in Th1-like Tregs treated with FTY720 compared to Th1-like Tregs, supporting the hypothesis that FTY720 treatment directly inhibits the generation of Th1-like Tregs (Fig. 1D, 1E).

**Figure 1.**
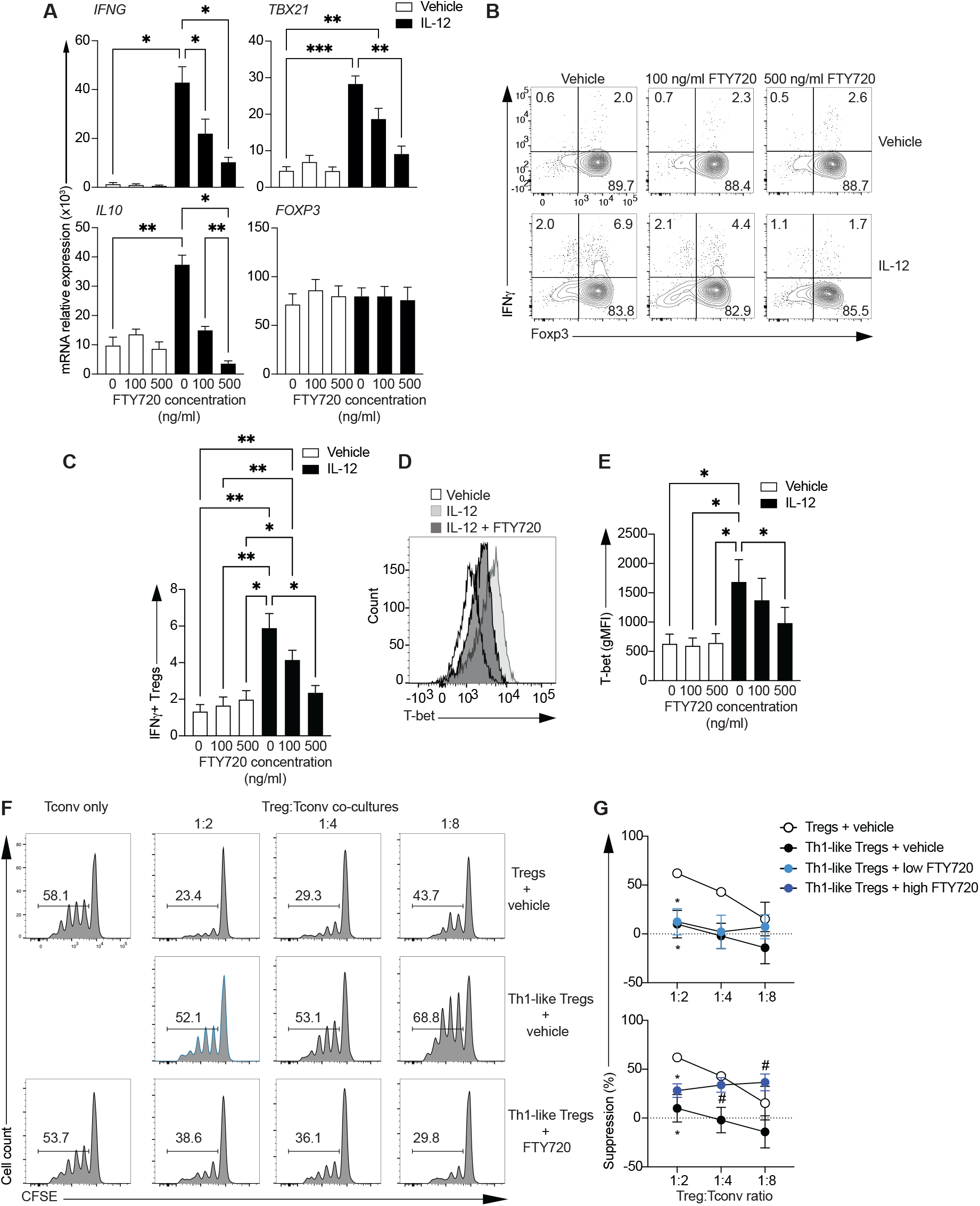
FTY720 inhibits the generation of Th1-like Tregs *in vitro*. Sorted Tregs from healthy individuals were stimulated with anti-CD3, anti-CD28 and IL-2 in the presence or absence of IL-12 and increasing doses of FTY720. **A**. Gene expression measured 48 hours after activation (n=4-9). Representative example (**B**) and summary (**C**) of IFN_γ_ production 4 days after initial activation and a 4 hour incubation with PMA and ionomycin in the presence of GolgiStop™ (n=8). Representative histogram (**D**) and summary (**E**) of T-bet expression measured 48 hours after activation and represented as geometric mean fluorescence intensity (gMFI, n=7). **F** and **G**. Sorted Tregs (CD4^+^CD25^high^CD127^low^) were stimulated as above in the presence or absence of IL-12. 72 hours later, cells were collected, washed and co-cultured with Treg-depleted, CFSE-labeled CD4^+^ T cells (Tconv) for 3.5 days in the presence of various concentrations of FTY720. **F**. Representative histogram of Tconv proliferation in co-culture at various Treg:Tconv ratios. **G**. Summary of percentage of suppression by Tregs and Th1-like Tregs in the presence or absence of FTY720 at various Treg:Tconv ratios (n=5). One-way ANOVA or mixed effects model with Tukey’s correction for multiple comparisons for **A, C** and **E**. Two-way ANOVA with Tukey’s correction for multiple comparisons in **G**. *p<0.05, **p<0.01, ***p<0.005.

To determine whether the inhibition of Th1-like Treg phenotype by FTY720 was accompanied by restored Treg function, Th1-like Tregs were generated *in vitro* with IL-12 and further co-cultured with CFSE-labeled conventional T cells (Treg-depleted CD4^+^ T cells, Tconv) in the presence or absence of FTY720. Tconv proliferation by CFSE dilution was measured as a readout for Treg suppressive capacity (Fig. 1F). As expected^2, 3^, *in vitro* generated Th1-like Tregs displayed impaired suppressive capacity as compared to Tregs, and this was restored by FTY720 to levels comparable to control Tregs (Fig. 1G). These data suggest that inhibition of S1P signaling by FTY720 inhibits the generation and impaired function of Th1-like Tregs.

### S1P signaling inhibition suppresses the activation of mTORC1

We went on to examine potential mechanisms underlying the inhibition of Th1-like Tregs by FTY720. We and others had previously shown that the PI3K/AKT/FOXO pathway is activated in Th1-like Tregs, and its inhibition restores Treg phenotype and function^2, 9^. Therefore, we examined the activation of AKT as a readout for PI3K signaling in Tregs and Th1-like Tregs in the presence or absence of FTY720 (Fig. 2). Th1-like Tregs displayed an increased phosphorylation of AKT at Thr 308 and Ser 473, the two residues that are required for full activation of AKT^36^ (Fig. 2A, 2B). Interestingly, FTY720 significantly inhibited Thr 308 phosphorylation of AKT, while not affecting phosphorylation of Ser 473 (Fig. 2B). FOXO1A and FOXO3A are downstream targets of PI3K/AKT and are involved in the generation of Th1-like Tregs^2, 37^. However, FTY720 did not inhibit the phosphorylation of FOXO1A or FOXO3A in Th1-like Tregs (Fig. 2C, 2D), suggesting that FTY720 is targeting other AKT downstream targets.

**Figure 2.**
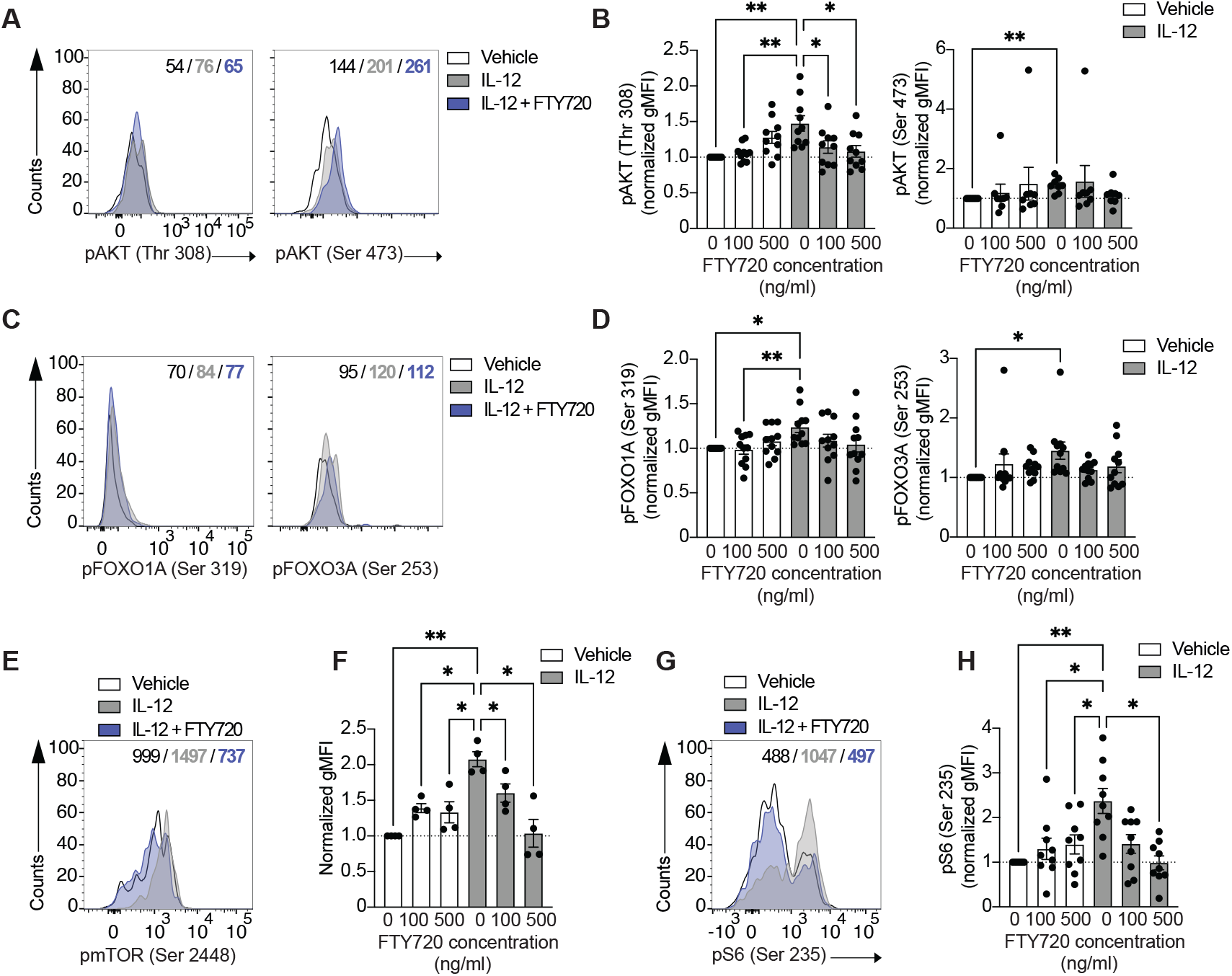
FTY720 inhibits mTORC1 signaling to suppress Th1-like Treg generation. CD4^+^ T cells from healthy individuals were stimulated with anti-CD3, anti-CD28 and IL-2 in the presence or absence of IL-12 and increasing doses of FTY720 for 18 hours and fixed for cellular staining. Representative histogram (**A**) and summary (**B**) of phosphorylation of AKT measured at Thr 308 (left, n=10) and Ser 473 (right, n=8). Representative histogram (**C**) and summary (**D**) of phosphorylation of FOXO1A (left, n=11) and FOXO3A (right, n=11). Representative histogram (**E**) and summary (**F**) of phosphorylation of mTOR (n=4). Representative histogram (**G**) and summary (**H**) of S6 phosphorylation (n=9). All summary plots are shown as normalized gMFI (to vehicle-treated Treg values). One-way ANOVA with Tukey’s correction for multiple comparisons for **B, D, F** and **H**. p<0.05, **p<0.01, ***p<0.005.

We decided to examine mTORC1 activation because it is downstream of PI3K/AKT signaling and has been implicated in controlling regulatory T cell function^38^. Interestingly, dysfunctional Th1-like Tregs displayed an increased phosphorylation of mTOR (Fig 2E) that was significantly suppressed with FYT720 in a dose-dependent manner (Fig 2F). In addition, phosphorylation of S6, a downstream target of mTORC1, was significantly increased in Th1-like Tregs as compared to Tregs (Fig. 2G), and FTY720 significantly decreased its activation to control levels (Fig. 2H).

mTORC1 has been shown to act as a negative and positive regulator of Treg development and function, and contrasting works suggest that its functions are associated with a fine tuning of its activity in different contexts^38^. We decided to examine whether inhibition of mTORC1 was sufficient to inhibit Th1-like generation (Fig. 3). For this, we stimulated Tregs and Th1-like Tregs in the presence of increasing concentrations of rapamycin, an mTORC1 inhibitor. Th1-like Tregs significantly decreased the expression of *IFNG* and *TBX21* in the presence of rapamycin in a dose-response manner (Fig. 3A), while no changes were observed in *FOXP3* gene expression. These results were confirmed at the protein level, with *in vitro*-generated Th1-like Tregs decreasing the production of IFN_γ_ in the presence of rapamycin (Fig. 3B, 3C) while maintaining the levels of FOXP3 expression (Fig. 3D). In agreement with the decrease in IFN_γ_ production, T-bet expression was significantly diminished in Th1-like Tregs in the presence of rapamycin (Fig. 3E, 3F), overall suggesting that mTORC1 inhibition is sufficient to inhibit Th1-like Treg generation. Mechanistically, rapamycin did not affect the activation of AKT (Thr 308) observed in Th1-like Tregs (Fig. 3G, 3H), but it significantly decreased the activation of mTOR (Fig. 3I, 3J) and its downstream target S6 (Fig. 3K, 3L).

**Figure 3.**
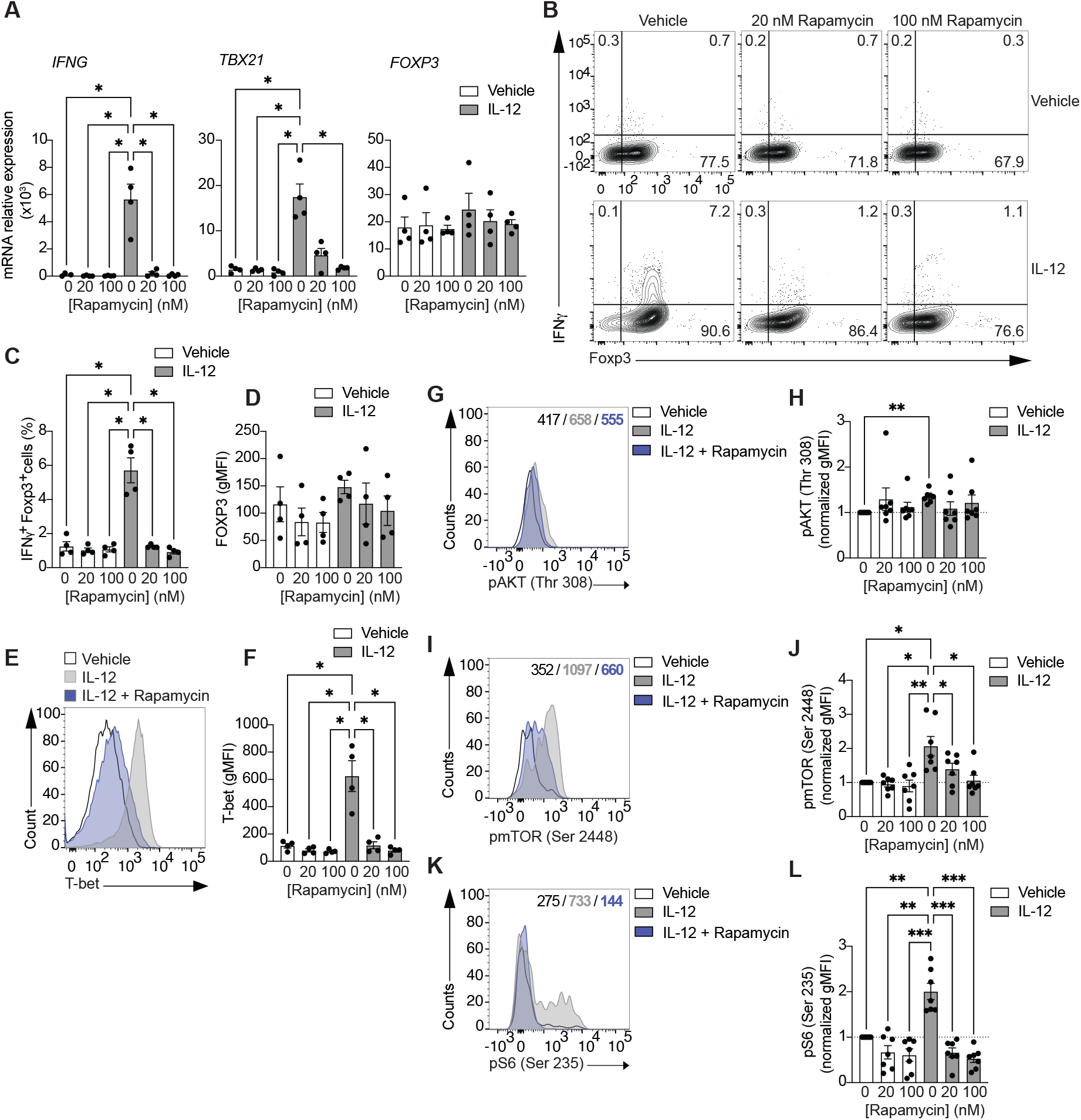
mTORC1 inhibition with rapamycin is sufficient to inhibit Th1-like Treg reprogramming. Sorted Tregs from healthy individuals were stimulated with anti-CD3, anti-CD28 and IL-2 in the presence or absence of IL-12 and increasing doses of rapamycin. **A**. Gene expression measured 48 hours after activation (n=4). Representative example (**B**) and summary (**C**) of IFN_γ_ production 4 days after initial activation and a 4 hour stimulation with PMA and ionomycin in the presence of GolgiStop™ (n=4). **D**. Summary of FOXP3 expression 48 hours after activation (n=4). Representative histogram (**E**) and summary (**F**) of T-bet expression measured 48 hours after activation and represented as geometric mean fluorescence intensity (gMFI, n=4). Representative histogram (**G**) and summary (**H**) of phosphorylation of AKT measured at Thr 308 (left, n=7). Representative histogram (**I**) and summary (**J**) of phosphorylation of mTOR (n=7). Representative histogram (**K**) and summary (**L**) of S6 phosphorylation (n=7). All summary plots are shown as normalized gMFI (to vehicle-treated Treg values). One-way ANOVA with Tukey’s correction for multiple comparisons for **A, C, D, F, H, J** and **L**. Only comparisons of Th1-like Tregs treated with vehicle to all other groups shown. p<0.05, **p<0.01, ***p<0.005.

Overall, these data suggest that FTY720 inhibits Th1-like Treg generation by suppressing the PI3K/AKT/mTORC1 pathway, and inhibition of mTORC1 by rapamycin is sufficient to suppress IL-12-driven Th1-like Treg generation *in vitro*.

### Th1-like Tregs undergo mitochondrial uncoupling that is reversed by S1P signaling inhibition

mTORC1 is an important nutrient sensor that integrates metabolic cues with signaling pathways. It has been shown to control mitochondrial biogenesis and activity, glycolysis and it facilitates the rewiring towards anabolic metabolism required for the activation and expansion of T cells^39^. Mitochondrial function is essential for Treg suppressive capacity^20, 21^, and therefore, we decided to explore if mitochondrial metabolism is altered in Th1-like Tregs, and whether S1P signaling inhibition suppresses Th1-like Treg generation by modulating it.

We used SCENITH™ to examine the metabolic phenotype of Th1-like Tregs and the effect of FTY720 in their dependence on specific metabolic pathways. SCENITH™ uses protein translation measured as puromycin incorporation as a readout for ATP production^40^ and determines the dependence of cells on mitochondrial respiration or glycolysis based on the use of inhibitors that specifically suppress those pathways. Th1-like Tregs did not display differences in their energetic status measured as puromycin incorporation as compared to Tregs, and FTY720 did not have any effect either (Fig. 4A). However, Th1-like Tregs significantly decreased their mitochondrial dependence, suggesting that there is a decrease in the proportion of ATP production that is dependent on oxidative phosphorylation (OXPHOS). FTY720 significantly restored this impairment to levels comparable to those of control Tregs (Fig. 4B). The decrease in mitochondrial dependence was confirmed using Seahorse assays (Fig. 4C). Basal oxygen consumption rate (OCR) was significantly decreased in *in vitro*-generated Th1-like Tregs as compared to control Tregs and FTY720 treatment of Th1-like Tregs restored it. Moreover, after inhibition of ATP synthase with oligomycin to determine the portion of basal respiration that is used to generate ATP (Fig. 4D), a decrease in the ATP produced by the mitochondria was observed in Th1-like Tregs, which was further restored to control Treg levels by FTY720 (Fig. 4D). Interestingly, no differences in glucose dependence (proportion of ATP production that is dependent on glucose oxidation) were observed in Th1-like Tregs (Fig. 4E) compared to control Tregs, and accordingly, basal extracellular acidification rate (ECAR) was similar in control and Th1-like Tregs, with FTY720 treatment not having a significant effect (Fig. 4F). However, Th1-like Tregs displayed a significant increase in their glycolytic capacity (maximum capacity to sustain ATP production when mitochondrial respiration is inhibited) relative to control Tregs and this was inhibited by FTY720 treatment (Fig. 4G). No changes in the overall capacity to use fatty acids and amino acids as fuels for ATP production when glucose oxidation is inhibited were observed in Th1-like Tregs.

**Figure 4.**
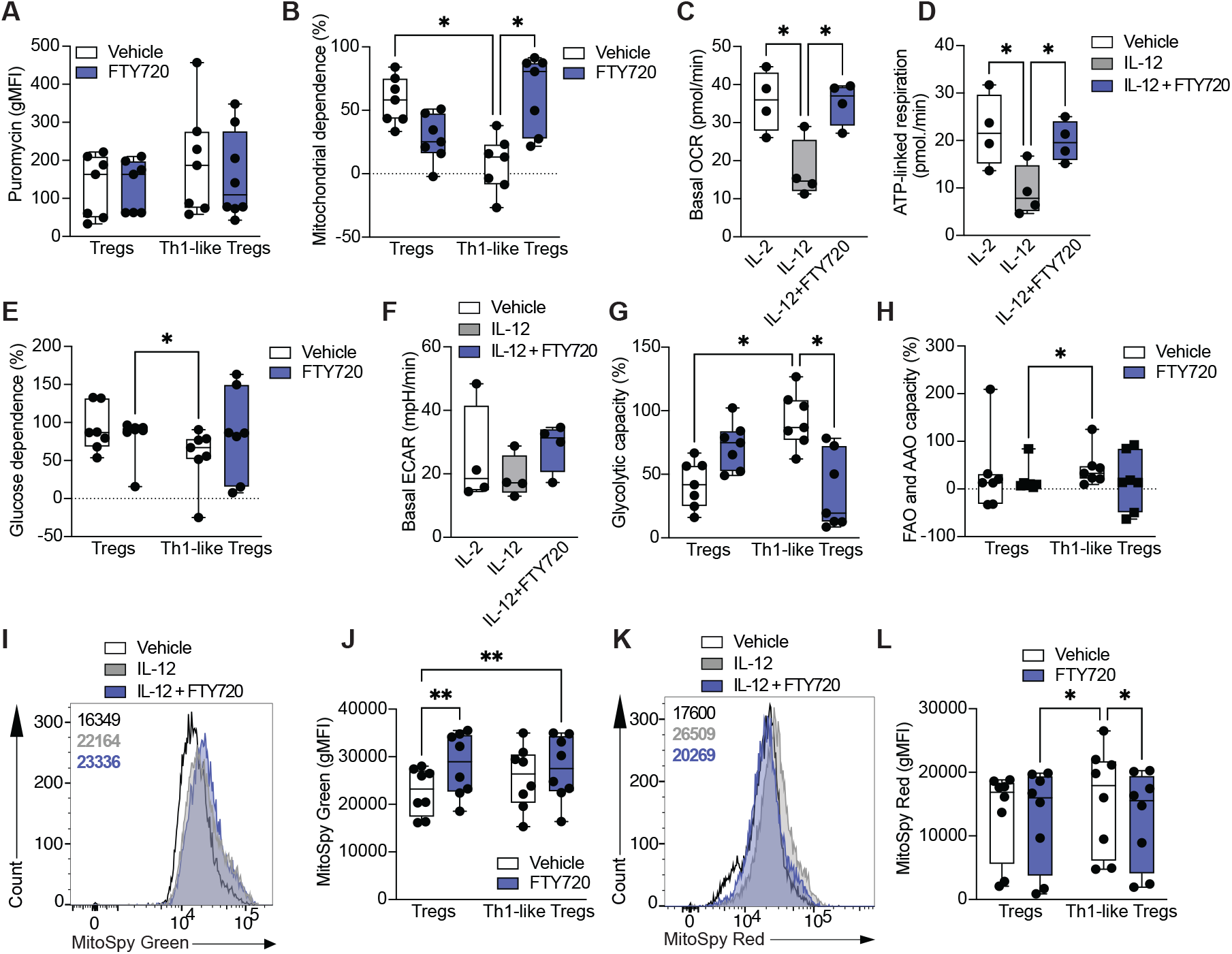
Th1-like Tregs display mitochondrial uncoupling that is reversed by S1P signaling inhibition. Sorted Tregs from healthy individuals were stimulated with anti-CD3, anti-CD28 and IL-2 in the presence or absence of IL-12 and FTY720 for 48 hours. **A**. Summary of puromycin incorporation measured as gMFI (n=7). **B**. Summary of percentage of mitochondrial dependence (n=7). **C**. Summary of basal oxygen consumption rate (OCR, n=4). **D**. Summary of ATP-linked respiration (n=4). **E**. Summary of percentage of glucose dependence (n=7). **F**. Summary of basal extracellular acidification rate (n=4). **G**. Summary of percentage of glycolytic capacity (n=7). **H**. Summary of percentage of fatty acid and amino acid oxidation (n=7). Representative histogram (**I**) and summary (**J**) of mitochondrial mass measured by MitoSpy Green incorporation (gMFI, n=8). Representative histogram (**K**) and summary (**L**) of mitochondrial membrane polarization measured by MitoSpy Red incorporation (gMFI, n=8). One-way ANOVA with Tukey’s correction for multiple comparisons for **C, D** and **F**. Two-way ANOVA with Tukey’s correction for multiple comparisons for **A, B, E, G, H, J** and L. p<0.05, **p<0.01.

To investigate the impairment in mitochondrial respiration in Th1-like Tregs in more detail, we measured mitochondrial mass (Fig. 4I, 4J) and mitochondrial membrane potential (Fig. 4K, 4L)^41^. We observed no significant changes in mitochondrial mass in Th1-like Tregs compared to control Tregs (Fig. 4I), but FTY720 increased mitochondrial mass in both Tregs and Th1-like Tregs (Fig. 4J). Unexpectedly, mitochondrial membrane potential was not impaired in Th1-like Tregs compared to control Tregs, but was slightly decreased by FTY720 treatment.

These results suggest that Th1-like Tregs undergo mitochondrial uncoupling characterized by a dissociation between mitochondria respiration-dependent ATP production and mitochondrial membrane potential. S1P signaling inhibition by FTY720 rebalances mitochondrial metabolism of Th1-like Tregs by increasing mitochondrial dependence and decreasing mitochondrial polarization.

### FTY720 rebalances Th1-like Treg mitochondrial metabolism in vivo

Patients with RRMS have an increased frequency of Th1-like Tregs *in vivo*^3^. Therefore, in order to determine whether FTY720 also led to a modification of mitochondrial function *in vivo*, we carried out a cross sectional study examining Tregs from healthy individuals and patients with RRMS either untreated or treated with FTY720 (Table 1). In agreement with our previous data^32^, untreated RRMS patients displayed increased levels of IFN_γ_-producing Tregs and higher expression of T-bet compared to Tregs isolated from healthy individuals *ex vivo* (Fig. 5A). FTY720 treatment inhibited Th1-like Tregs *in vivo*, and Tregs isolated from FTY720-treated RRMS patients produced significantly less IFN_γ_ and downregulated T-bet expression, while no changes were observed in FOXP3 and IL-10 protein levels (Fig. 5A). Moreover, *in vivo* FTY720 treatment inhibited the aberrant AKT activation observed in Tregs from untreated RRMS patients compared to healthy individuals (Fig. 5B), but no significant changes were observed in FOXO3A phosphorylation in Tregs from FTY720-treated compared to untreated RRMS patients (Fig. 5C), in agreement with the *in vitro* data (Figure 2). We went on to determine the activation status of mTORC1 (Fig. 5D, 5E). mTOR phosphorylation showed a trend towards increased expression in untreated RRMS patients, and FTY720 treatment significantly reduced it. This was accompanied by an increased phosphorylation of S6 in untreated RRMS patients compared to healthy individuals. FTY720-treated patient Tregs displayed a trend towards decreased pS6 as compared to untreated RRMS patients. When exploring the metabolic phenotype of Tregs from the three groups of patients, we found that FTY720-treated RRMS patients showed a significant increase in mitochondrial dependence and a decrease in glycolytic capacity compared to untreated RRMS patients, in agreement with the data generated *in vitro* (Fig. 5F, 5G). Finally, in order to determine whether the extent of metabolic alterations observed in RRMS patients correlated with the presence of Th1-like Tregs, we tested whether mitochondrial changes, i.e. mitochondrial dependence and glycolytic capacity correlated with T-bet expression as a readout for the size of the Th1-like Treg population. Mitochondrial dependence showed a negative correlation with T-bet expression in untreated RRMS patients, suggesting that Tregs from those untreated RRMS patients with increased Th1-like phenotype also display lower mitochondrial dependence. This correlation was lost in patients treated with FTY720 (Fig. 5H). Similarly, glycolytic capacity positively correlated with T-bet expression only in untreated RRMS patients (Fig. 5I).

**Figure 5.**
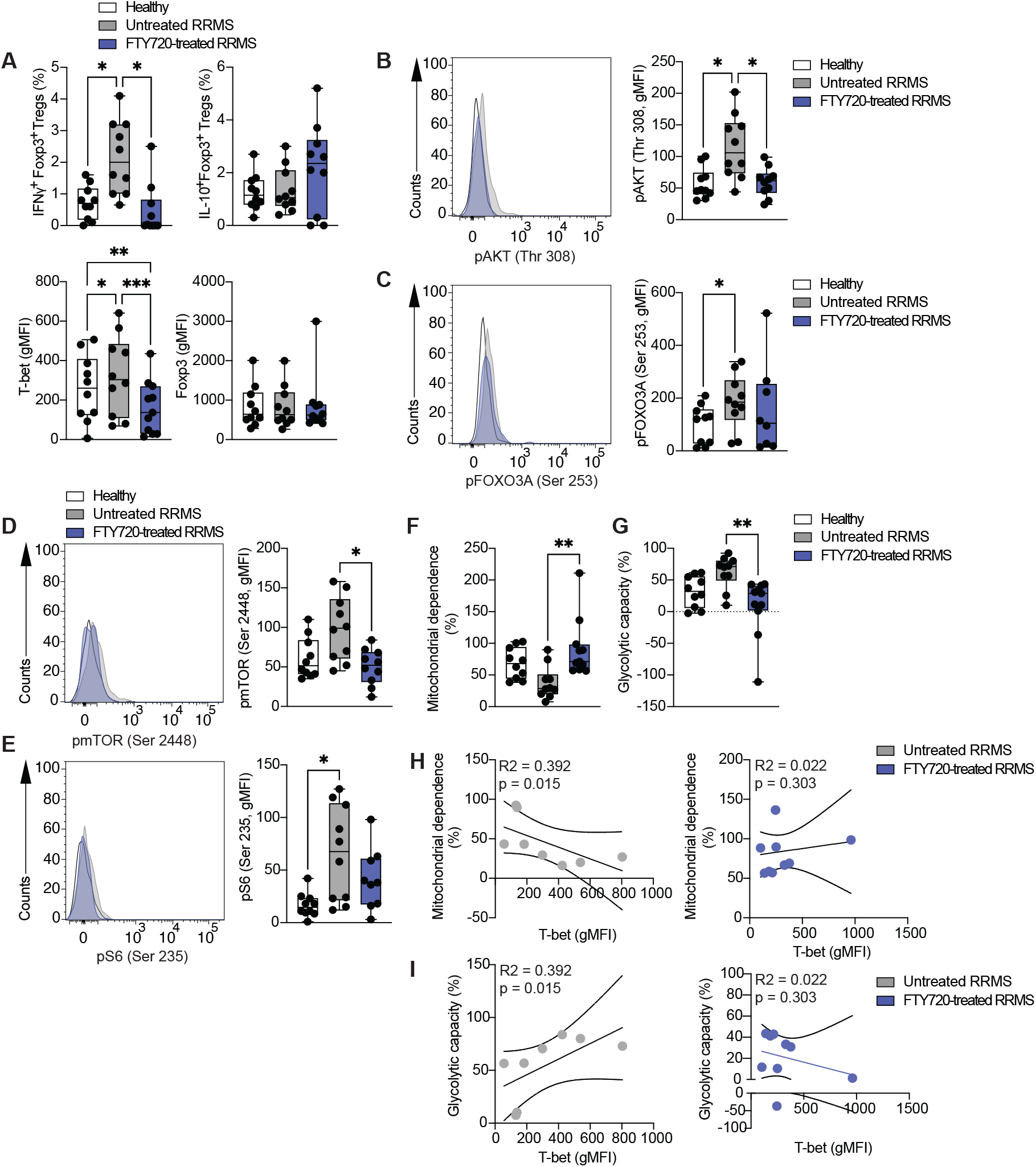
*In vivo* FTY720 treatment restores mitochondrial imbalance in Tregs from RRMS patients. Sorted Tregs from healthy individuals (white) and patients with RRMS either untreated (grey) or treated with FTY720 (blue) were stimulated with PMA and ionomycin in the presence of GolgiStop™ for 4 hours (**A**) or left unstimulated (**B**-**G**). **A**. Summary of percentage of IFN_γ_ and IL-10 production and T-bet and FOXP3 expression (gMFI, n=10-11). **B**. Representative histogram (left) and summary (right) of phosphorylation of AKT measured at Thr 308 (n=10). **C**. Representative histogram (left) and summary (right) of phosphorylation of FOXO3A (n=8-10). **D**. Representative histogram (left) and summary (right) of mTOR phosphorylation (n=10). **E**. Representative histogram (left) and summary (right) of S6 phosphorylation (n=9-10). **F**. Summary of percentage of mitochondrial dependence (n=10-11). **G**. Summary of percentage of glycolytic capacity (n=10-11). **H**. Correlation of mitochondrial dependence (percentage) with T-bet expression (gMFI) in untreated (left) and FTY720-treated (right) RRMS patients. **I**. Correlation of glycolytic capacity (percentage) with T-bet expression (gMFI) in untreated (left) and FTY720-treated (right) RRMS patients. One-way ANOVA with Tukey’s correction for multiple comparisons for **A, C, D, E, F** and **G**. Spearman correlation for **H** and **I**. p<0.05, **p<0.01, ***p<0.005.

These data suggest that *in vivo* FTY720 treatment inhibits the Th1-like phenotype in RRMS patients by targeting the PI3K/AKT/mTORC1 pathway and rebalancing mitochondrial metabolism to reverse mitochondrial uncoupling.

## Discussion

In this work we demonstrate that S1P signaling can modulate human Treg stability and function *in vitro* and *in vivo*, as its inhibition with FTY720 prevents the generation of a dysfunctional and pro-inflammatory Th1-like Treg phenotype by inhibiting mTORC1 signaling and reversing Th1-like Treg mitochondrial uncoupling.

The main reported role of S1P signaling in immune cells is controlling lymphocyte migration. However, increasing evidence suggests that S1P signaling is involved in other migration-independent cellular processes in immune cells^24^. Moreover, sphingolipid metabolism has been found to be dysregulated in many human diseases, including cancer, inflammation, atherosclerosis and asthma^42^, with therapies targeting S1P signaling being effective at controlling disease in MS patients^25, 26^. Thus, *in vivo* S1P signaling inhibition with FTY720 treatment in RRMS patients results in increased frequencies of Tregs but a pronounced depletion of effector T cells in the periphery, with changes in Treg phenotype suggestive of increased function^32^. These data support the hypothesis that S1P signaling regulates Teff and Treg migration differently, but most importantly, that S1P signaling is involved in other cellular functions besides migration. The cell type-specific expression of S1PR as well as their differential coupling to various G proteins likely contribute to the diverse functional outcomes of S1P signaling^43^. In this regard, our observations are in agreement with early findings in rodents that S1PR1 overexpression exerts a cell-autonomous negative regulation on mouse Treg function^29^.

Our data suggest that Th1-like Treg generation requires mTORC1 activation, which is inhibited by FTY720 treatment. mTORC1 is an important nutrient sensor and metabolic regulator of glucose and mitochondrial metabolism that is activated downstream of AKT. The involvement of mTORC1 in the control of Treg phenotype and function is complex and contrasting results have been reported regarding its role in inhibiting or promoting Treg development and function. These works have demonstrated that mTORC1 is required for functional competency of thymic-derived Tregs and have highlighted the delicate control mTORC1 activity is subjected to in T cells and Tregs^38, 44^. In our experimental setting, inhibition of mTORC1 by either FYT720 or rapamycin led to an impairment in the generation of Th1-like Tregs and restoration of their suppressive function. In agreement with these data, overactivation of mTORC1 by Treg-specific deletion of TSC1 has been shown to cause a decrease in Treg suppressive capacity and acquisition of an effector-like phenotype in a colitis model^45^.

Mitochondrial uncoupling, the functional dissociation between the generation of mitochondrial membrane potential and its use for ATP synthesis, can be the consequence of multiple processes including proton leak, inducible leak through uncoupler proteins (UCPs) and electron leak^46^. While initially thought to be detrimental for mitochondrial function, the identification of uncoupling proteins (UCP-1 to 5 in humans) suggests that mitochondrial uncoupling is involved in other physiological processes^46^. For example, the expression of some uncoupling proteins such as UCP-2 have been shown to be induced by antigen stimulation of CD8^+^ T cells, suggesting that mitochondrial uncoupling is part of the physiological response to antigen^47^. In addition, polyclonal stimulation of both CD4^+^ and CD8^+^ T cells *in vitro* increases UCP-2 expression^48^. Little is known about how mitochondrial uncoupling controls the function of immune cells, and what endogenous pathways control mitochondrial uncoupling in immune cells. Our results suggest that mitochondrial uncoupling is involved in the generation of dysfunctional Th1-like Tregs and potentially favored by the S1P-mTORC1 signaling axis, as inhibition with FTY720 restores the mitochondrial metabolic imbalance *in vitro* and *in vivo*. Questions remain related to the specific mechanism that S1P signaling triggers to favor mitochondrial uncoupling during Th1-like Treg reprogramming and the contribution of mTORC1 signaling.

Increasing evidence associates mitochondrial dysfunction to a number of inflammatory, autoimmune and chronic conditions^49, 50, 51, 52^. In the case of RRMS patients, our data are in agreement with works demonstrating that T cells and Tregs from treatment naïve patients show decreased glycolysis and oxygen consumption rate *ex vivo*^53, 54^, suggesting that mitochondrial targeting in RRMS and potentially in other autoimmune conditions that present with similar Treg phenotypes^55^, could be considered as a potential novel therapeutic target.

## Methods

### Patient sample and clinical data collection

Patients and healthy donors who met the eligibility criteria were recruited and provided informed consent (ethics approval obtained from the South Central - Berkshire Research Ethics Committee, REC reference number 20/SC/0308). Peripheral blood was collected from all participants and processed following a common standard operating protocol (Table 1).

### Cell culture reagents and antibodies

Cells were cultured in RPMI 1640 supplemented with 10 mM HEPES Buffer Sodium, 0.05DmM non-essential amino acids, 1 mM sodium pyruvate, 2 mM L-Glutamine, 100 U/mL and 100 μg/mL penicillin/streptomycin respectively (all from GIBCO), and 5% human heat-inactivated AB serum (Sigma). The antibodies used for stimulation were anti-human CD3 (clone UCHT1, plate bound) and anti-human CD28 (clone 28.2, BD Biosciences, San Jose, CA) at 1 μg/mL. Treg Inspector Beads (Miltenyi Biotec, Bergisch Gladbach, Germany) were used in suppression assays following manufacturer’s recommendations. FTY720 (Sigma-Aldrich) was used at 100 and 500 ng/mL. Rapamycin (Sigma-Aldrich) was used at 20 and 100 nM. IL-2 was obtained through the AIDS Research and Reference Reagent Program, Division of AIDS, National Institute of Allergy and Infectious Diseases (NIAID), National Institutes of Health (NIH) and was used at 50 U/mL. The antibodies used in this work are summarized in Table 2.

### PBMC isolation, storage, and thawing

Peripheral blood mononuclear cells (PBMCs) were isolated by Ficoll-Paque PLUS (Cytiva) gradient centrifugation less than 2 hours after blood collection. The PBMC layer was collected, washed with PBS, resuspended at 20 million cells/mL in fetal bovine serum supplemented with 10% DMSO and stored at -150 °C.

For PBMC thawing, vials were thawed in a pre-warmed water bath at 37 °C, transferred to 15 mL conical tube and 6 mL of pre-warmed complete media was added. The tubes were subsequently centrifuged for 10 min at 300 x *g* and resuspended in warm complete media at 5 million cells/mL.

### Flow cytometry staining for regulatory T cell immunophenotyping

PBMCs were thawed and CD4^+^ T cells were isolated by immunomagnetic negative selection (EasySep™ Human CD4^+^ T Cell Enrichment Kit, StemCell Technologies). 100,000-200,000 CD4^+^ T cells per well were plated in 96-well V-bottom plates and rested for 1.5 hours at 37 °C, 5% CO_2_ in complete media. For *ex vivo* phenotypic characterization, the cells were stained immediately after resting according to the protocol described hereafter, with the antibodies detailed in Table 2. For the characterization of the cytokine profile, the cells were incubated for 4 hours in 200 μL of complete media with 50 ng/mL PMA and 250 ng/mL ionomycin in the presence of GolgiStop™ and stained immediately after with antibodies detailed in Table 2.

All centrifugation steps in this protocol were carried out at 300 x *g* for 10 minutes. First, cells were stained with LIVE/DEAD Fixable Blue Dead Cell Dye (Thermo Fisher Scientific) according to the manufacturer’s protocol. A Fc receptor (FcR) blocking step with FcR Blocking Reagent Human (Miltenyi Biotec) was performed with the cell surface antibody staining. The cells were subsequently fixed with the Foxp3/Transcription Factor Staining Buffer Set (Thermo Fisher Scientific) according to the manufacturer’s protocol. Where relevant, an additional step for intracellular staining was added after fixation, using the FoxP3 staining buffer kit. The cells were then washed and resuspended in 250 μL of PBS.

The samples were run on a Fortessa instrument (BD Biosciences) and analyzed using FlowJo v10.0 (BD Biosciences). Tregs were gated on size and granularity (lymphocyte gate), single cells, live cells, CD3^+^CD4^+^CD25^high^CD127^low^.

### In vitro Th1-like Treg generation

Fresh PBMC were isolated and CD4^+^ T cells were enriched by immunomagnetic negative selection as described above. CD4^+^ T cells were stained with antibodies to CD4, CD25 and CD127 and Tregs (CD4^+^CD25^high^CD127^low^) and conventional T cells (Tconv, Treg-depleted CD4^+^ T cells) were sorted on a FACS Aria II. Sorted Treg cells were plated in 96-well U-bottom plates (50,000 cells/well) pre-coated with 1 μg/mL plate-bound anti-CD3 (clone UCHT1), and stimulated in complete media supplemented with soluble anti-CD28 (1 μg/mL), with 50 U/mL IL-2 and with or without 20 ng/mL IL-12 for 4 days at 37 °C, 5% CO_2_. Some wells also received FTY720 besides IL-12 or rapamycin. Cells were lysed 48 hours after stimulation for gene expression analysis. After 4 days in culture, cells were restimulated with 50 ng/mL PMA and 250 ng/mL ionomycin in the presence of GolgiStop™ for the characterization of the cytokine profile.

### Metabolic profiling using SCENITH™

SCENITH™ is a flow cytometry-based assay for profiling cell energy metabolism with single cell resolution^40^. Briefly, rapid changes in protein translation upon metabolic pathway inhibition are monitored using puromycin incorporation to newly synthesized proteins as a reliable readout for protein synthesis, which is tightly coupled to ATP production and therefore can be used as a readout for the impact of metabolic pathway inhibition on the energetic status of the cells.

PBMCs were thawed and plated at 300,000-500,000 cells per well in 96-well V-bottom plates and rested for 1.5 hours at 37 °C, 5% CO_2_ in complete media. The cells were then treated for 45 minutes at 37 °C, 5% CO_2_ with vehicle (Control, Co), 100 mM 2-deoxy-D-glucose (DG, Sigma-Aldrich), 1 μM oligomycin (O, Sigma-Aldrich) or a combination of both drugs (DGO). 10 μg/mL puromycin was added to all conditions. Subsequently, cells were pelleted by centrifugation for 7 minutes at 400 x *g* and stained according to the published protocol^40, 56^. Cells were washed, resuspended in 250 μL of PBS and run on a Fortessa instrument (BD Biosciences). Data were analyzed using FlowJo v10.0 (BD Biosciences). Tregs were gated based on size and granularity, singlets, live cells, CD3^+^CD4^+^CD25^high^CD127^low^.

For the analysis of the energetic status of the cells, puromycin geometric mean fluorescence intensity was analyzed in each of the four conditions mentioned above (Co, DG, O, DGO). The percentage of glucose dependence was calculated using this formula: 100*(Co-DG)/(Co-DGO); and the percentage of mitochondrial dependence was calculated as 100*(Co-O)/(Co-DGO). Finally, glycolytic capacity (%) was defined as 100-mitochondrial dependence, and fatty acid and amino acid oxidation capacity (%) was defined as 100-glucose dependence.

### Quantification of mRNA expression by real-time PCR

RNA was isolated using the RNeasy Micro Plus Kit (Qiagen) following the manufacturer’s guidelines in Appendix D of the Qiagen RNeasy handbook. Isolated RNA was converted to complementary DNA by reverse transcription (RT) with random hexamers and Multiscribe RT (TaqMan Reverse Transcription Reagents, Thermo Fisher Scientific). The reactions were set up using the manufacturer’s guidelines and run on a QS5 Studio Real-Time PCR instrument (Thermo Fisher Scientific). Values are represented as the difference in cycle threshold (Ct) values normalised to *B2M* expression for each sample as per the following formula: Relative RNA expression = (2-ΔCt) x 10^3^.

### Flow cytometry-based phosphorylation assays

For phosphorylation assays with patient samples (Figure 5), PBMCs were thawed and CD4^+^ T cells were isolated by immunomagnetic negative selection (EasySep™ Human CD4^+^ T Cell Enrichment Kit, StemCell Technologies). 250,000-500,000 CD4^+^ T cells per well were plated in 96-well V-bottom plates, rested for 2 hours at 37 °C, 5% CO_2_ in complete media, and fixed immediately after. For phosphorylation assays of Th1-like Tregs, PBMC were isolated and CD4^+^ T cells were isolated fresh as above. Subsequently, 250,000-500,000 CD4^+^ T cells per well were plated in 96-well U-bottom plates pre-coated with 1 μg/mL plate-bound anti-CD3 (clone UCHT1), and stimulated in complete media supplemented with soluble anti-CD28 (1μg/mL), with 50 U/mL IL-2 and with or without 20 ng/mL IL-12 for 18 hours at 37°C, 5% CO_2_. Some wells also received FTY720 besides IL-12. 18 hours later, the cells were fixed following the protocol described hereafter.

For all conditions, cells were fixed in pre-warmed Fix Buffer I (BD Phosflow) for 20 minutes at 37 °C, 5% CO_2_ and permeabilized with ice-cold Perm Buffer III (BD Phosflow) overnight at -20 °C. Cells were subsequently stained in PBS for 1 hour at room temperature with the antibodies detailed in Table 2. Subsequently, cells were washed in PBS and resuspended in 250 μL PBS. The samples were run on a Fortessa instrument (BD Biosciences) and analyzed using FlowJo v10.0 (BD Biosciences). Tregs were gated on size and granularity, singlets, live cells, CD25^high^FoxP3^+^.

### Suppression assays

PBMC were thawed and stained with antibodies recognizing CD3, CD4, CD25 and CD127. Tregs (CD3^+^CD4^+^CD25^high^CD127^low^) and Tresp (CD3^+^CD4^+^CD25^-/int^CD127^+^) were sorted using a FACSAria™ Fusion flow cytometer (BD Biosciences). Propidium Iodide was added right before the sort to exclude dead cells.

Subsequently, Tconv were stained with CellTrace™ CFSE, and a suppression assay was set up in a 96-well round-bottom plate with 5 different conditions: Tconv only (0:1), co-cultures of Tconv and Tregs at 3 different Treg:Tconv ratios (1:2, 1:4, 1:8); and Tconv without stimulation as a negative control for proliferation. Treg Suppression Inspector beads (Miltenyi Biotec) were added to all conditions (except Tconv without stimulation) at a 2:1 ratio to the cells. The cells were cultured in 200 μL complete media for 3.5 days and were then stained according to the protocol described for Treg immunophenotyping, with antibodies to CD4, CD25 and Ki67. The samples were run on a Fortessa instrument (BD Biosciences) and analyzed using FlowJo v10.0 (BD Biosciences). Tconv were gated on size and granularity, singlets, live cells, CD4^+^ and CFSE^+^. Suppressive capacity (%) was calculated using this formula: 100-((Co-culture division/Tconv only division)*100).

### Statistical analysis

Data were analyzed using GraphPad Prism version 10. Normal distribution of the data was tested using the Anderson-Darling and D’Agostino and Pearson normality tests, or Shapiro-Wilk test for those datasets with a small number of replicates. Normally distributed data by at least one of the two tests was analyzed by one- or two-way ANOVA when comparing more than two groups of one or two independent variables, respectively. A two-tailed t-test was used to compare two groups. Data were expressed as mean ± s.e.m. Where data are presented as box and whiskers, the boxes extend from the 25^th^ to the 75^th^ percentile and the whiskers are drawn down to the minimum and up to the maximum values. Horizontal lines within the boxes denote the median. p values >0.05 were considered statistically significant.

## Acknowledgements

We thank Ms Lolena Parreno and the rest of the MS clinical team at Charing Cross Hospital for patient recruitment and sample obtention. We thank Dr Isabel Correa-Otero and Ms Radhika Patel for their help with flow cytometry sorting and members of the MDV lab for critical reading of the manuscript. This work was funded by a BBSRC grant (BB/W001055/1) to MDV.

## Author contribution

RC performed experiments, analyzed the data, and wrote the manuscript; NG and CS performed experiments and analyzed the data; NN, AAM and AKM performed experiments; RA provided anti-puromycin antibody and advised on SCENITH™ data generation and analysis; AS, RN and JV recruited RRMS patients and provided advice on the clinical aspects of the work; MDV designed the study, analyzed data, wrote the manuscript, and obtained funding. All authors revised and contributed to the editing of the manuscript.

## Supplementary Figures

**Supplementary Figure 1. FTY720 treatment does not affect Treg viability**. Sorted Tregs from healthy individuals were stimulated with anti-CD3, anti-CD28 and IL-2 in the presence or absence of IL-12 and FTY720 for 4 days and viability was examined after 4 hours of PMA and ionomycin stimulation. Representative dot plots (**A**) and summary (**B**) of percentage of live Tregs. One-way ANOVA with Tukey’s correction for multiple comparisons.

**Supplementary Figure 2. FTY720 treatment does not alter FOXP3 expression**. Sorted Tregs from healthy individuals were stimulated with anti-CD3, anti-CD28 and IL-2 in the presence or absence of IL-12 and FTY720 for 4 days and FOXP3 expression was examined after 4 hours of PMA and ionomycin stimulation. Summary of n=8 experiments. One-way ANOVA with Tukey’s correction for multiple comparisons.

## Tables

**Table 1. Patient characteristics.**

**Table 2. Antibodies used in this work.**

## Notes

### Competing Interest Statement

The authors have declared no competing interest.

