## Supplementary material for "Sphingosine-1-phosphate signaling inhibition suppresses Th1-like Treg generation by reversing mitochondrial uncoupling": Tables 1 and 2: Table 1.docx

|  | Fingolimod | Untreated |
| --- | --- | --- |
| Total number | 12 | 10 |
| Age on collection | | |
| Mean (SD) | 49 (8) | 39 (9) |
| Minimum-maximum | 37-58 | 26-57 |
| Sex | | |
| Male | 3 (25%) | 2 (20%) |
| Female | 9 (75%) | 8 (80%) |
| Ethnicity | | |
| White (caucasian) | 8 (67%) | 5 (50%) |
| White(other) | 1 (8%) | 2 (20%) |
| Asian (other) | 2 (17%) | 2 (20%) |
| Black/African/Caribbean | 1 (8%) | 1 (10%) |
| Age on diagnosis | | |
| Mean (SD) | 37 (9) | 38 (9) |
| Minimum-maximum | 23-54 | 26-54 |

**Table 1. Patient characteristics.**
