## Supplementary material for "Sphingosine-1-phosphate signaling inhibition suppresses Th1-like Treg generation by reversing mitochondrial uncoupling": Tables 1 and 2: Table 2.docx

| **Assay** | **Target** | **Clone** | **Fluorochrome** | **Manufacturer** | **Catalog #** |
| --- | --- | --- | --- | --- | --- |
| Treg phenotyping, SCENITH | CD3 |  | BV785 | Biolegend | 300471 |
| Treg phenotyping,  SCENITH | CD25 |  | PE | Biolegend | 356104 |
| Treg phenotyping,  SCENITH | CD127 |  | FITC | BD Phamingen | 560549 |
| Treg phenotyping | T-bet |  | PE-Cy7 | Invitrogen | 25-5825-82 |
| Treg phenotyping | Foxp3 |  | AF700 | eBioscience | 56-4776-41 |
| Treg phenotyping | IL-10 |  | APC | BD Pharmingen | 554707 |
| Treg phenotyping | IFNγ |  | BV605 | Biolegend | 505839 |
| SCENITH | CD4 |  | V500 | BD Horizon | 560769 |
| SCENITH | CD45RO |  | BV421 | BD Horizon | 562641 |
| SCENITH | T-bet |  | PerCP-Cy5.5 | Invitrogen | 25-5825-82 |
| SCENITH | FoxP3 |  | PE-Cy7 | eBioscience | 45-4776-73 |
| SCENITH | Puromycin |  | AF647 | Gift from Dr Argüello | N/A |
| Phosflow | FoxP3 |  | eFluor450 | eBioscience | 48-4777-42 |
| Phosflow | CD25 |  | AF700 | BD Pharmingen | 561398 |
| Phosflow | FoxO3a (pS253) |  | AF488 | Bioss | bs-3140R-A488 |
| Phosflow | S6 (pS235/S236) |  | AF647 | BD Biosciences | 56043 |
| Phosflow | AKT (pS473) |  | PE | BD Biosciences | 560378 |
| Phosflow | mTOR (pS2448) |  | AF647 | BD Biosciences | 564242 |
| Phosflow | AKT (pT308) |  | PE | BD Biosciences | 558275 |

**Table 2. Antibodies used in this work.**
