## Supplementary figures and images for "Sphingosine-1-phosphate signaling inhibition suppresses Th1-like Treg generation by reversing mitochondrial uncoupling"

### Supplementary Figures

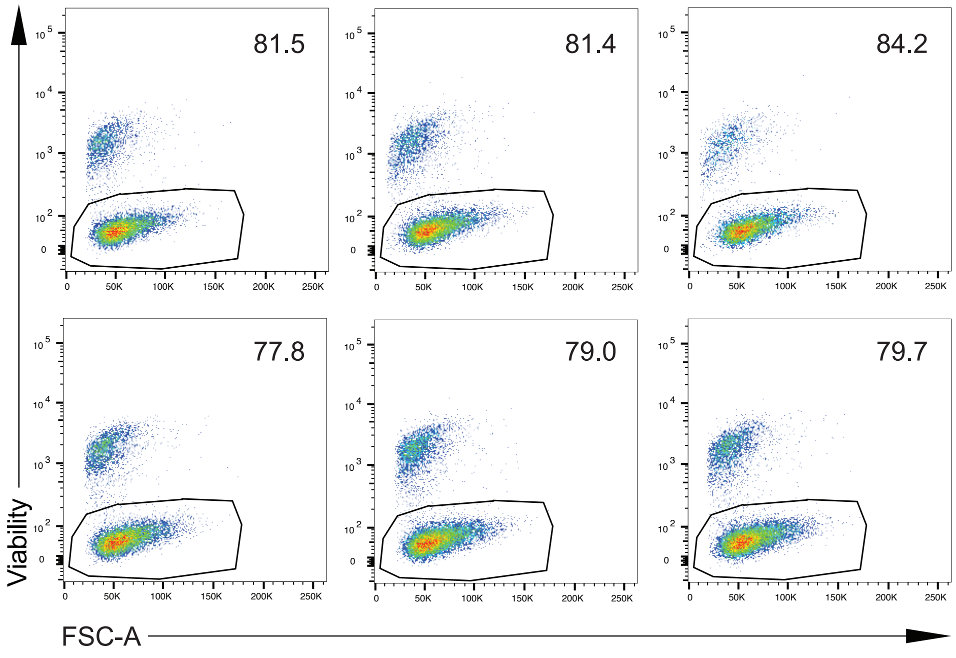


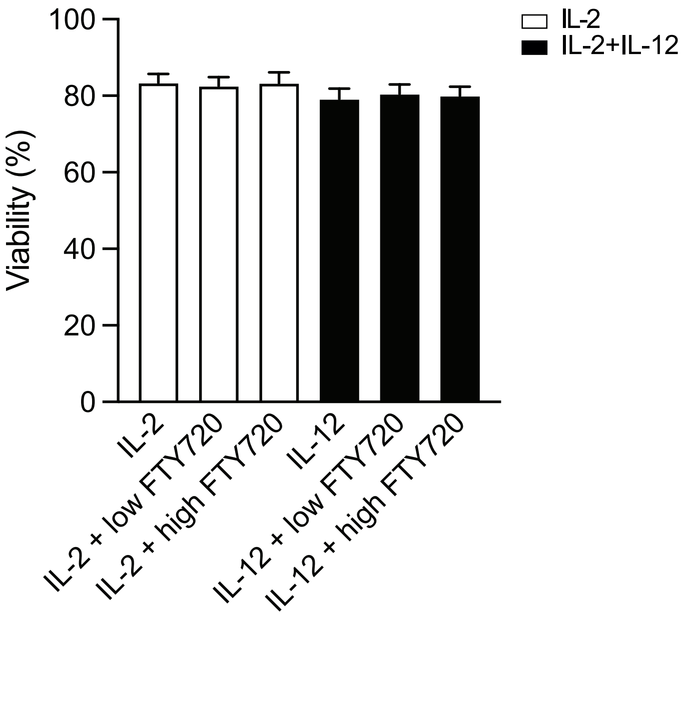


Supplementary Figure 1.

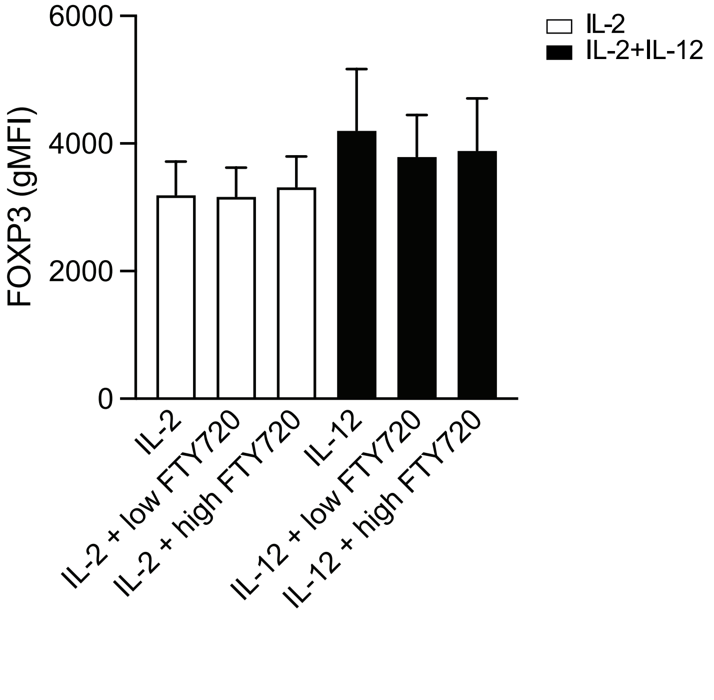


Supplementary Figure 2
